# Predatory plants and patchy cows: modeling cattle interactions with toxic larkspur amid variable heterogeneity

**DOI:** 10.1101/487561

**Authors:** Kevin E. Jablonski, Randall B. Boone, Paul J. Meiman

## Abstract

The most common explanations for the evolution and persistence of herd behavior in large herbivores relate to decreased risk of predation. However, poisonous plants such as larkspur (*Delphinium* spp.) can present a threat comparable to predation. In the western United States, larkspur diminishes the economic and ecological sustainability of cattle production by killing valuable animals and restricting management options. Recommendations for mitigating losses have long focused on seasonal avoidance of pastures with larkspur, despite little evidence that this is practical or effective. Our ongoing research points to the cattle herd itself as the potential solution to this seemingly intractable challenge and suggests that larkspur and forage patchiness may drive deaths. In this paper, we present an agent-based model that incorporates neutral landscape models to assess the interaction between plant patchiness and herd behavior within the context of poisonous plants as predator and cattle as prey. The simulation results indicate that larkspur patchiness is indeed a driver of toxicosis and that highly cohesive herds can greatly reduce the risk of death in even the most dangerous circumstances. By placing the results in context with existing theories about the utility of herds, we demonstrate that grouping in large herbivores can be an adaptive response to patchily distributed poisonous plants. Lastly, our results hold significant management-relevant insight, both for cattle producers managing grazing in larkspur habitat and in general as a call to reconsider the manifold benefits of herd behavior among domestic herbivores.

## Introduction

Of the more than 60 species of larkspur (*Delphinium* spp. L.) found in North America, at least eleven are known to cause significant cattle losses, primarily those species found on rangelands in the western United States and Canada (Green et al. 2009, Welch et al. 2015a). High levels of norditerpinoid alkaloids, which cause neuromuscular paralysis when consumed in sufficient quantity, are the chief culprit in these toxicosis deaths (Ralphs et al. 1988, Manners et al. 1995). Total yearly deaths due to larkspur toxicosis have been estimated at 2-5% of grazing cattle in some regions, with an annual cost of $234 million to producers (Pfister et al. 1997, Knight and Walter 2001, Welch et al. 2015a). This makes larkspur one of the leading causes of death losses in the US cattle industry (Knight and Walter 2001).

Grazing management recommendations in larkspur habitat have long focused on seasonal avoidance, aimed at reducing exposure during spring and early summer when alkaloid concentration is highest (Pfister et al. 1997, Welch et al. 2015a). This strategy creates problems of its own as producers lose flexibility to meet their management objectives, both economic and ecological, with little evidence of reduced losses. Because of this, many producers appear to simply accept the risk of deaths, achieving gains when lucky and losses when not. One alternative to avoidance is to manage grazing such that no individual is able to consume a lethal dose of alkaloids, regardless of season. Our recent paper (Jablonski et al. 2018) presented an agent-based model that indicated this may be possible if cattle are managed for high stocking density, high herd cohesion, or both.

While our findings in Jablonski et al. (2018) were relevant to grazing management within the habitat of a particular larkspur species (*Delphinium geyeri Green*), the results also pointed towards interesting relationships between plant patchiness, herd behavior, and toxicosis that we explore further here. Specifically, we used modeling to test two general hypotheses, that: (1) larkspur patchiness drives alkaloid toxicosis deaths, and (2) overlap between larkspur and desirable forage drives alkaloid toxicosis deaths. We explore both hypotheses within the context of variations in herd cohesion, using data from *D. geyeri*, wherein *N*-(methylsuccinimido)-anthranoyllycoctonine (MSAL) type alkaloids are the dominant toxin (Panter et al. 2002).

### Neutral landscape models

A test of the influence of larkspur patchiness and larkspur-forage overlap on toxicosis required a model with variable landscapes, rather than the realistic but static landscape of Jablonski et al. (2018). Specifically, this meant separating larkspur and forage distribution from one another and varying patchiness while maintaining a realistic landscape with respect to cattle grazing. For this, we used neutral landscape models, which are the most common landscape modelling approach used in ecological studies, with frequent application to habitat fragmentation, animal movement models, and metapopulation analysis (Gardner and Urban 2007, Synes et al. 2016). With a primary aim of improving understanding of how ecological processes are affected by spatial structure, neutral landscape models are ideal for testing the consequences of varying spatial heterogeneity on foraging outcomes (With and King 1997). However, we are unaware of previous application of neutral landscape models to cattle grazing dynamics.

### Behavioral ecology of herds

Important context for this study comes from the literature on grouping in large herbivores, where behavioral ecologists continue to debate the evolution and utility of herd behavior (e.g. Makin et al. 2017, Ireland and Ruxton 2017, Stutz et al. 2018). The most widely studied explanations for herd behavior relate to decreased risk of predation (Davies et al. 2012, Ebensperger and Hayes 2016). Of particular relevance to the cattle-larkspur interaction is what Krause and Ruxton (2002) call dilution. Dilution refers to a 1/N effect whereby an attacking predator can only capture a limited number of prey at a time and individual risk therefore declines with increasing group size. Important considerations for the dilution effect are variation in the likelihood of being attacked among individuals and the relative conspicuousness of larger versus smaller groups (Krause and Ruxton 2002).

A second relevant mechanism for decreased predation risk in herds is predator avoidance, also known as encounter dilution. In this case, predators with limited perceptual range encounter clumped prey at a lower frequency than single prey (Krause and Ruxton 2002). It is necessary to consider dilution and encounter dilution in context with one another, as increased detectability can offset the benefits of herd members’ reduced likelihood of death when encountering a predator (Turner and Pitcher 1986).

We examine larkspur as predator and cattle as prey. This is a novel approach, and poisonous plants certainly differ from typical predators in many ways. However, there is enough similarity to enable this “plants as predators” concept to be useful addressing both theoretical and practical questions.

### Agent-based modeling

Agent-based models are computational simulation tools that focus on bottom-up encoding of individual “agent” behaviors as they interact with one another and the environment (Grimm 1999, McLane et al. 2011). Agent-based models are particularly useful in modeling complex systems where the results of interactions between system elements are not easily predicted, and thus useful for simulating the behavior of social herbivores foraging in a heterogeneous environment (Dumont and Hill 2004, Grimm et al. 2005). Nevertheless, they have thus far been little used in improving our understanding of livestock behavior and management.

In this paper, we present an agent-based model simulation of cattle grazing with varied herd cohesion in larkspur-rich pastures with varied plant patchiness. Our approach represents a novel application of neutral landscape models and agent-based models to the relationship between herbivore grazing behavior and environmental heterogeneity. The results offer insights to landscape ecology, behavioral ecology, and livestock grazing management, and point toward a fundamental reconsideration of the importance of herd behavior among domestic herbivores.

## Methods

### Overview

The model functions as a mechanistic effects model (Grimm and Martin 2013) whereby cattle seek to maximize forage intake within behavioral and physiological bounds and are exposed to toxic alkaloids via consumption of larkspur distributed within the forage. Deaths are a product of temporal intensity of larkspur consumption with passing time as a mitigating factor via metabolism. The guiding principles of model design were behavior-based encoding (McLane et al. 2011) of cattle activities, based in the literature and our own livestock management experience, and parsimony aimed at including only those behaviors and landscape variables relevant to the question at hand. Model evaluation followed the process of “evaludation” laid out by Augusiak et al. (2014).

Cows are classified as leaders (5% of herd), followers (85%), or independents (10%), with leaders making decisions about broad-scale movements away from relatively over-grazed areas (known as site changes) and independents being less tied to the herd than the other cows (Sato 1982, Harris et al. 2007). Other than seeking drinking water and making site changes, all cow movements in the model are aimed at moving closer to herdmates and/or maximizing the amount of forage in the next grazing location, depending on desired herd proximity. Consumption of forage occurs in line with standard rates from the literature (Laca et al. 1994, WallisDeVries et al. 1998). Forage and larkspur amounts decrease when eaten and do not regrow within the model run, which is equivalent to 18 days.

Other details of model function can be found in the complete Overview, Design Concepts, and Details (ODD; Grimm et al. 2010) description in Jablonski et al. (2018). Here we focus on model elements that have changed, using the ODD format but omitting sections where methods were the same.

### Purpose

The agent-based model tests the effect of co-varying herd cohesion (also known as troop length; Shiyomi and Tsuiki 1999), larkspur patchiness, and larkspur-forage overlap on cases of lethal alkaloid toxicosis caused by larkspur similar in size and toxicity to measured values for *D. geyeri*. We developed and executed the model in NetLogo 6.01, using the BehaviorSpace tool to implement simulations (Wilensky 1999).

### Entities and state variables

The model has two kinds of entities: pixels representing 1 m^2^ of land and agents representing 500 kg adult cows (1.1 animal-units). Because computational demands would be higher with additional covariates, and spatial extent was expected to be minimally influential, we shrank the model landscape to ¼ the size of that of pasture 16 of the Colorado State University Research Foundation Maxwell Ranch on which the model in Jablonski et al. (2018) was based. This created a model landscape of 832 x 790 pixels (0.83 km x 0.79 km, equal to 65.73 ha), all of which are accessible to the cows. Note that, for clarity, we will refer to each 1 m^2^ land area as a pixel, rather than as a patch, the typical nomenclature for agent-based models. We use “patch” in the landscape ecology sense to refer to an area of habitat that is relatively discrete from its surroundings in relation to some phenomenon of interest (Turner and Gardner 2015).

Stocking density was set at 1.0 animal-units • ha^-1^ throughout the simulation, totaling 59 cows. Herd cohesion was determined using herd-distance-factor (HDF), in which increasing values indicate greater inter-animal distance. All other state variables, including role, MSAL-tolerance, and larkspur-attraction were assigned in the same way as Jablonski et al. (2018). All functionally relevant state variables for pixels and cows, as well as global variables and inputs, are described in Table 1.

**Table 1.**
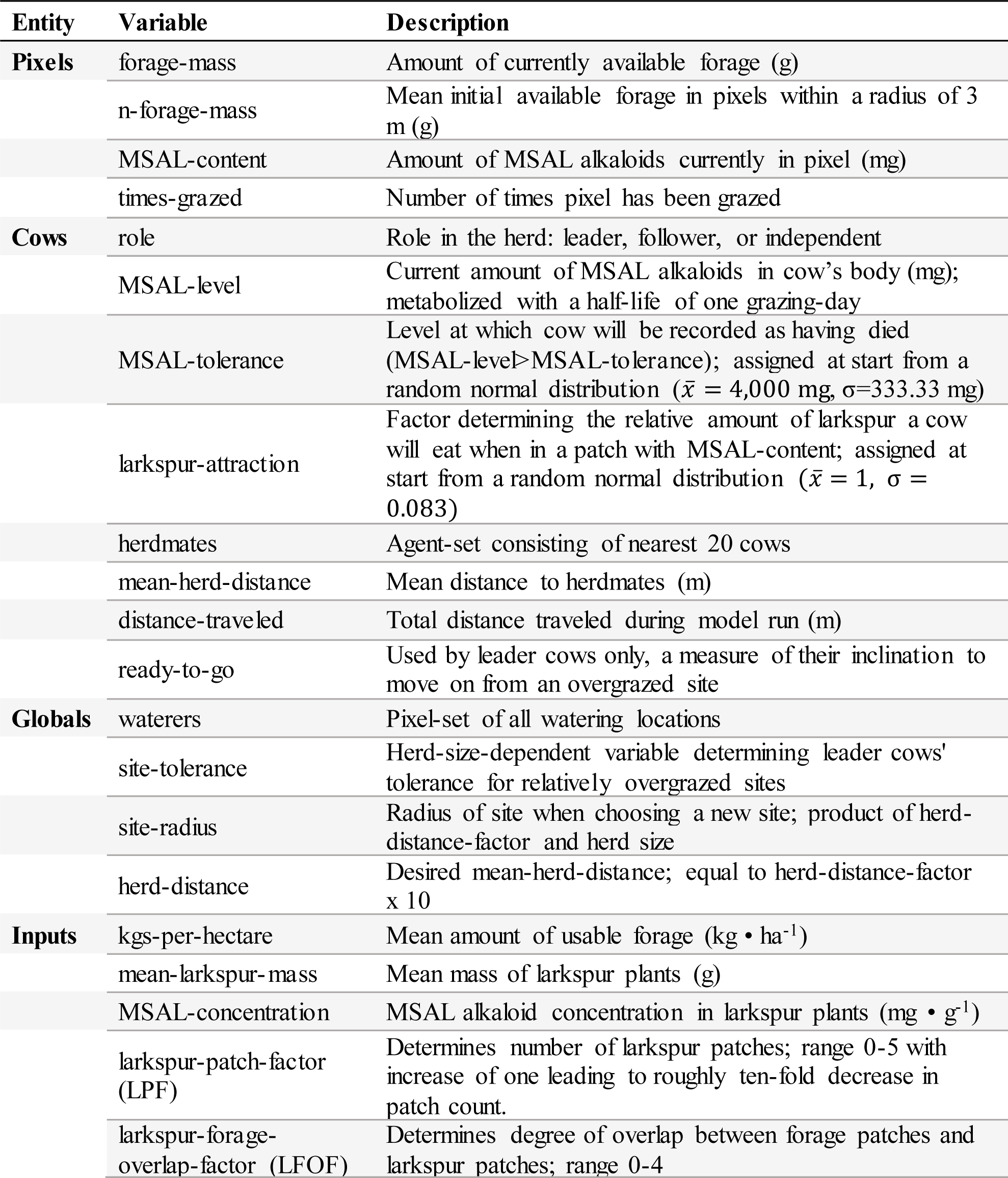
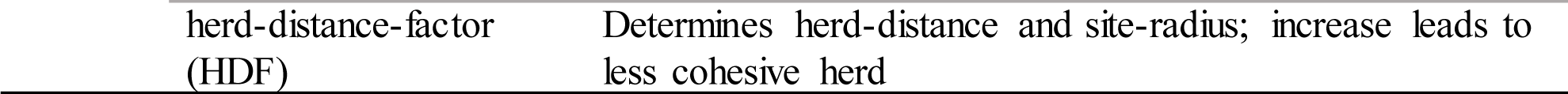
Relevant model variables. Sources for variable parameters are cited in the body of the text.

| Entity | Variable | Description |
| --- | --- | --- |
| **Pixels** | forage-mass | Amount of currently available forage (g) |
|  | n-forage-mass | Mean initial available forage in pixels within a radius of 3 m (g) |
|  | MSAL-content | Amount of MSAL alkaloids currently in pixel (mg) |
|  | times-grazed | Number of times pixel has been grazed |
| **Cows** | role | Role in the herd: leader, follower, or independent |
|  | MSAL-level | Current amount of MSAL alkaloids in cow's body (mg); metabolized with a half-life of one grazing-day |
| | MSAL-tolerance | Level at which cow will be recorded as having died (MSAL-level > MSAL-tolerance); assigned at start from a random normal distribution ($\bar{x} = 4,000$ mg, $\sigma = 333.33$ mg) |
| | larkspur-attraction | Factor determining the relative amount of larkspur a cow will eat when in a patch with MSAL-content; assigned at start from a random normal distribution ($\bar{x} = 1$, $\sigma = 0.083$) |
|  | herdmates | Agent-set consisting of nearest 20 cows |
|  | mean-herd-distance | Mean distance to herdmates (m) |
|  | distance-traveled | Total distance traveled during model run (m) |
|  | ready-to-go | Used by leader cows only, a measure of their inclination to move on from an overgrazed site |
| **Globals** | waterers | Pixel-set of all watering locations |
|  | site-tolerance | Herd-size-dependent variable determining leader cows' tolerance for relatively overgrazed sites |
|  | site-radius | Radius of site when choosing a new site; product of herd-distance-factor and herd size |
|  | herd-distance | Desired mean-herd-distance; equal to herd-distance-factor x 10 |
| **Inputs** | kgs-per-hectare | Mean amount of usable forage ($\text{kg} \cdot \text{ha}^{-1}$) |
|  | mean-larkspur-mass | Mean mass of larkspur plants (g) |
| | MSAL-concentration | MSAL alkaloid concentration in larkspur plants ($\text{mg} \cdot \text{g}^{-1}$) |
|  | larkspur-patch-factor (LPF) | Determines number of larkspur patches; range 0-5 with increase of one leading to roughly ten-fold decrease in patch count. |
|  | larkspur-forage-overlap-factor (LFOF) | Determines degree of overlap between forage patches and larkspur patches; range 0-4 |

|  |  |
| --- | --- |
| herd-distance-factor<br>(HDF) | Determines herd-distance and site-radius; increase leads to<br>less cohesive herd |

Note that death occurs when an individual cow exceeds its assigned value for MSAL-tolerance at the end of a grazing-day. However, the animal is not removed from the herd, but instead is recorded as having died, has its MSAL-level set to zero, and continues in the model. This preserves herd dynamics for the entire model run and makes it possible for total model-run deaths to exceed 59.

### Process overview and scheduling

Fig 1 illustrates the model execution process for each tick. Each cow moves through each step of the process, but only performs those steps linked to its role. Only elements that have changed from Jablonski et al. (2018) are described below, with explanation for the change.

**Fig 1.**
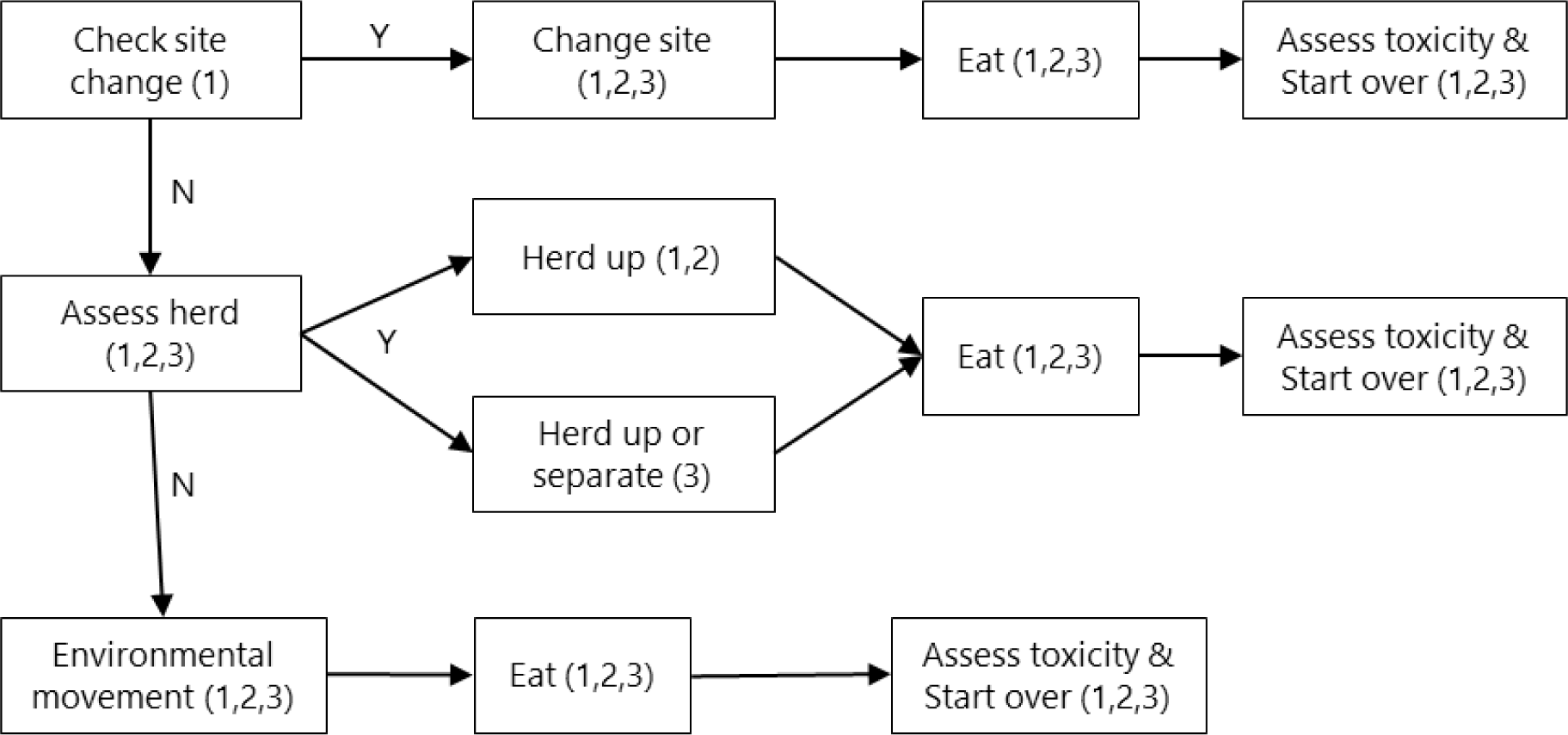
Pseudo-coded flow chart of model processes, with role of cows executing each process in parentheses. 1=leader, 2=follower, 3=independent.

Check hydration: This process is not found in Fig 1, as it was eliminated in favor of a single end-of-grazing-day water visit by all cows. Because hydration was previously linked to forage consumption, artificially high levels of forage heterogeneity necessitated a simplified water visit routine. One visit per day achieves this without otherwise sacrificing realistic model function. Assess herd: Herd-based movements are fundamentally the same as in Jablonski et al. (2018), with individuals moving closer to the herd centroid when mean-herd-distance exceeds herd-distance. However, we altered the minimum movement distance when “herding up” such that it is now based on the cow’s current distance from the herd centroid. This was to accommodate movement patterns in the tightly cohesive herds modeled at the low end of the herd-distance-factor range, where an arbitrary static minimum movement distance may cause them to frequently move through the herd and then beyond their desired mean-herd-distance, resulting in “ping-pong” type movements.

### Design concepts

#### Stochasticity

Distinct from Jablonski et al. (2018), the environment was highly stochastic between different levels of larkspur-patch-factor and larkspur-overlap factor and even within different iterations of identical values for those factors.

### Initialization

Input values for number of larkspur plants and forage mass within the modeled landscape were derived from the measured values from pasture 16 (Jablonski et al. 2018). This provides an input value of 107,500 total larkspur plants on the landscape. The model distributes these plants among pixels according to a Poisson distribution with a mean of 2.5 larkspur plants per square-meter pixel, resulting in 43,000 pixels with larkspur. This means that individual pixels with larkspur are equally likely to be dangerous regardless of their spatial arrangement, an essential condition for testing the effect of patchiness.

Landscape initialization within the model begins by using an input value for larkspur-patch-factor (LPF) to randomly locate *p* larkspur patch origins, with *p* = 43,000/10^*LPF*, rounded up to the nearest integer (i.e., 1 ≤ *p* ≤ 43,000) At each larkspur patch origin, a modified random walk is used to create realistic larkspur patch patterns. In this random walk, a temporary agent is created that visits each patch origin location. After placing a random Poisson-distribution-determined number of larkspur plants in the origin pixel, the agent then executes random turns and one-pixel steps, placing larkspur plants whenever landing on a pixel with zero currently present. This random walk proceeds in a given patch origin area until 43,000/*p* pixels have had larkspur plants placed in them. The agent then proceeds to the next patch origin location, following the same steps until all patch origin locations have been visited. Lastly, pixels are assigned an MSAL-content value based on larkspur plant count and input values for mean-larkspur-mass and MSAL-concentration.

Forage initialization begins with random placement of 80% of the total forage mass (100% being equal to ¼ of the forage mass in pasture 16) across the landscape, according to a normal distribution with a mean based on the input value for kgs-per-hectare. The remaining 20% is assigned according to the input value for larkspur-forage-overlap-factor (LFOF). For LFOF=0, all remaining forage is placed into forage patches (created using a similar random walk) that occupy 5% of the total land area. These forage patches do not overlap with larkspur patches. For LFOF=4, all remaining forage is placed within the larkspur patches and there are no forage patches. Values from 1-3 place increasing amounts of forage within the larkspur patches and decreasing amounts in the forage patches. We chose the values of 20% of forage in patches and 5% of land area in forage patches to approximate the forage heterogeneity found in pasture 16.

Instead of the seasonal stream watering locations found in pasture 16, the model places watering points in each corner and in the center of the landscape to ensure limited effect of distance from water (Bailey and Provenza 2008). Waterers are created as circular locations with a radius of 5 m. Finally, the model creates 59 cows (1.0 AU • ha^-1^) and places them at the central watering location. All other pixel values (Table 1) are derived from the various input values above.

### Simulation

We used the BehaviorSpace tool in Netlogo to run a full factorial simulation using eight levels of larkspur patchiness (LPF: 0, 1, 2, 3, 3.5, 4, 4.5, and 5), five levels of larkspur-forage overlap (LFOF: 0, 1, 2, 3, and 4), and six levels of herd cohesion (HDF: 0.5, 1, 2, 4, 8, and 16). With 30 replications of these 240 combinations, we executed 7,200 total model runs. The computational demands for this required creation and use of a virtual machine with 64 processors and 360 GB of RAM in Google Compute Engine (Google, Inc. 2018).

Input mean-larkspur-mass was 3.5 g and MSAL-concentration was 3.0 mg • g^-1^, representative of an excellent growing year with alkaloids at high levels. The input value for kgs-per-hectare was 500 kg • ha^-1^. Individual model run duration was 18 grazing-days, resulting in consumption of approximately 40% of available forage. All of these input values are based on our measurements from the Maxwell Ranch.

### Observation

As in Jablonski et al. (2018), data related to alkaloid intake were of prime importance, with deaths quantified according to a tolerance threshold (MSAL-tolerance) based on dose-response studies with larkspur (Welch et al. 2015b). The model also recorded numerous other data related to herd interactions, cow behavior, and landscape structure for purposes of model verification. These include inter-animal distance, frequency of herd-based movements, site-change frequency, travel distance, grazing impact, and mean larkspur count in pixels, among others.

In addition to model-run level outputs, each model run also recorded daily alkaloid consumption data for each cow. For 7,200 runs this amounted to 7.65 million data points. We compiled and organized this dataset using OpenRefine 3.0 (Google/Open source 2018). We also used this daily dataset to generate statistics on consumption for each individual grazing-day (n=129,600).

### Statistical analysis

To assess landscape structure, we analyzed a sample (n=10 for each level of larkspur-patch-factor) of the generated landscapes using class metrics in Fragstats 4.2.1 (McGarigal et al. 2012). We used the metrics number of patches (NP), percent land area in patches (PLAND; used to confirm uniformity), largest patch index (LPI), edge density (ED), clumpiness index (CLUMPY), and percent like adjacencies (PLADJ) (McGarigal et al. 2002).

We used both JMP 13.0.0 and R statistical software, version 3.5.1 for data exploration, analysis, and visualization (SAS Institute 2016, R Core Team 2018). We used the R base package to generate linear models, and the package MuMIn to compare models with AICc (Anderson 2008). We used the package ggplot2 in R to generate explanatory graphics.

## Results

### Model output verification

Because we have made only minor changes to grazing behavior in the model, we refer the reader to Jablonski et al. (2018) for results and discussion of output verification as it relates to cows. However, because landscape generation is greatly altered, we report landscape metrics in Table 2. Of the measured metrics, largest patch index and edge density were most strongly correlated with LPF. Fig 2 shows example landscapes at different combinations of LPF and LFOF.

**Table 2.** Mean landscape metrics for sample landscapes (n=10 per level) generated at different levels of larkspur-patch-factor (LPF). Reference McGarigal et al. (2002) for descriptions of metrics.

| LPF | Number<br>of<br>patches | Pct.<br>land in<br>patches | Largest<br>patch<br>index | Edge<br>density | Clumpiness<br>index | Pct. like<br>adjacencies |
| --- | --- | --- | --- | --- | --- | --- |
| 0 | 32463.10 | 6.54 | 0.00 | 2442.92 | 0.00 | 6.52 |
| 1 | 3400.10 | 6.54 | 0.01 | 1235.56 | 0.50 | 52.69 |
| 2 | 331.50 | 6.54 | 0.10 | 720.41 | 0.71 | 72.38 |
| 3 | 31.00 | 6.54 | 0.58 | 498.00 | 0.80 | 80.91 |
| 3.5 | 10.20 | 6.54 | 1.50 | 439.49 | 0.82 | 83.15 |
| 4 | 3.80 | 6.54 | 3.01 | 389.41 | 0.84 | 85.05 |
| 4.5 | 1.80 | 6.54 | 3.93 | 347.22 | 0.86 | 86.67 |
| 5 | 1.00 | 6.54 | 6.54 | 343.99 | 0.86 | 86.76 |

**Fig 2.**
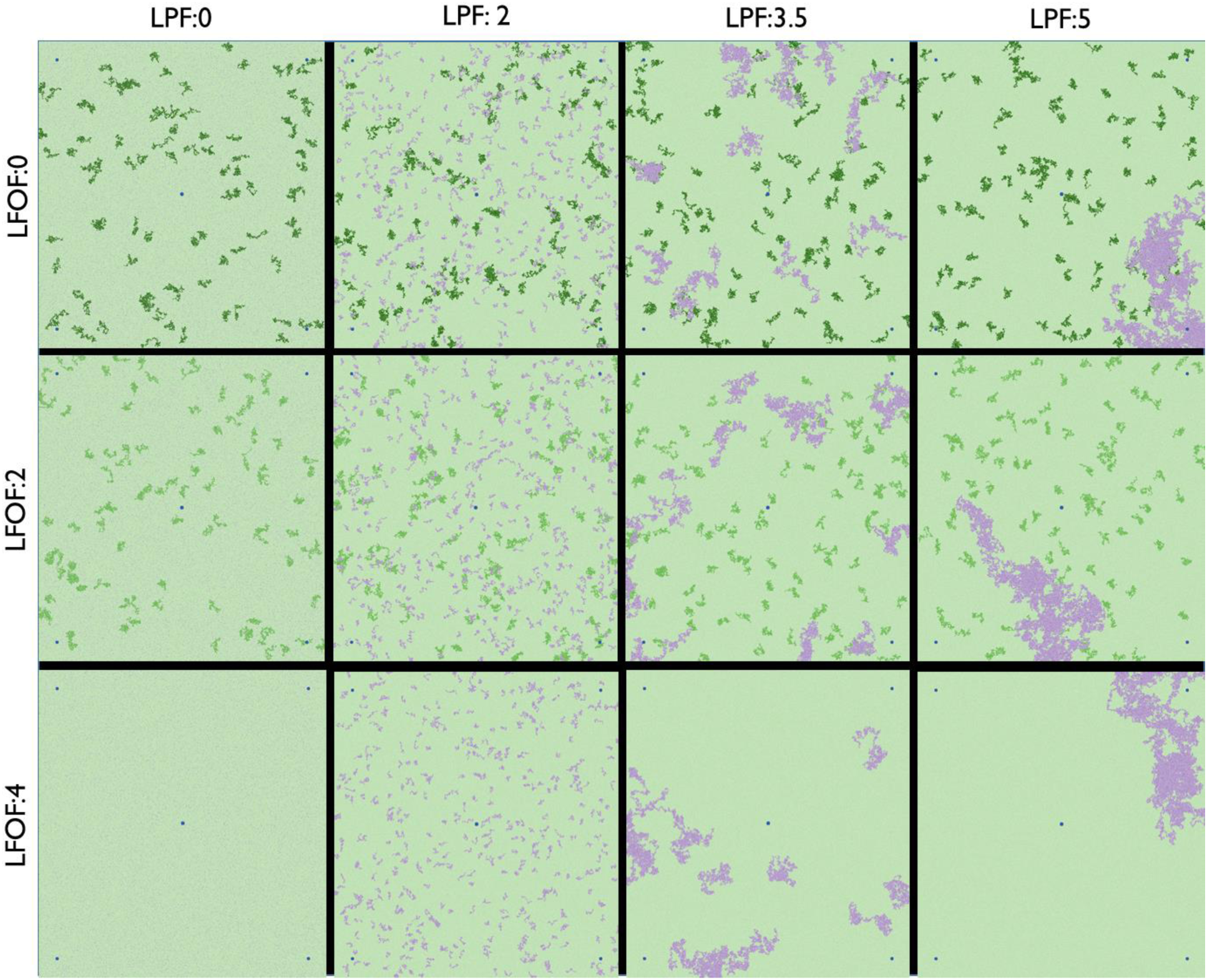
Sample landscapes for different levels of larkspur-patch-factor (LPF) and larkspur-forage-overlap (LFOF). Green indicates the distribution of forage, with darker green equal to more forage (forage patches), and purple indicates larkspur. No forage patches are visible at LFOF=4 because they are obscured by the larkspur. Watering locations are blue.

Note that, although HDF sets the desired maximum distance from herdmates (herd-distance), herds do not necessarily strictly adhere to this parameter. This is particularly true in less cohesive herds, where actual mean distance from herd mates was much lower than the maximum allowed by the HDF setting. For example, at HDF=16, herd-distance is set at 160 m, but overall mean distance from herdmates for all model runs at this level was 104.0 m, with a range from 83.6 m to 118.8 m. This is likely due to the overall size of the pasture and the time between regrouping at watering locations. Only at the lowest level of HDF (0.5) was overall mean herdmate distance at the maximum, as herdmates were essentially forced to stay closer to one another than foraging behavior would otherwise require.

### Toxicosis mechanism

In Jablonski et al. (2018) we identified the key mechanism for reducing larkspur deaths as narrowing the variation in larkspur consumption among individuals in the herd, with associated reduction in the count and extremity of outliers. As would be expected, deaths were once again strongly linked to this mechanism, with model-run standard deviation of daily alkaloid intake presenting a particularly striking pattern, wherein the likelihood and count of deaths increased once the standard deviation exceeded a threshold of 500 mg (Fig 3). Overall, at least one death occurred in 33.7% of model runs and on 6.2% of grazing-days.

**Fig 3.**
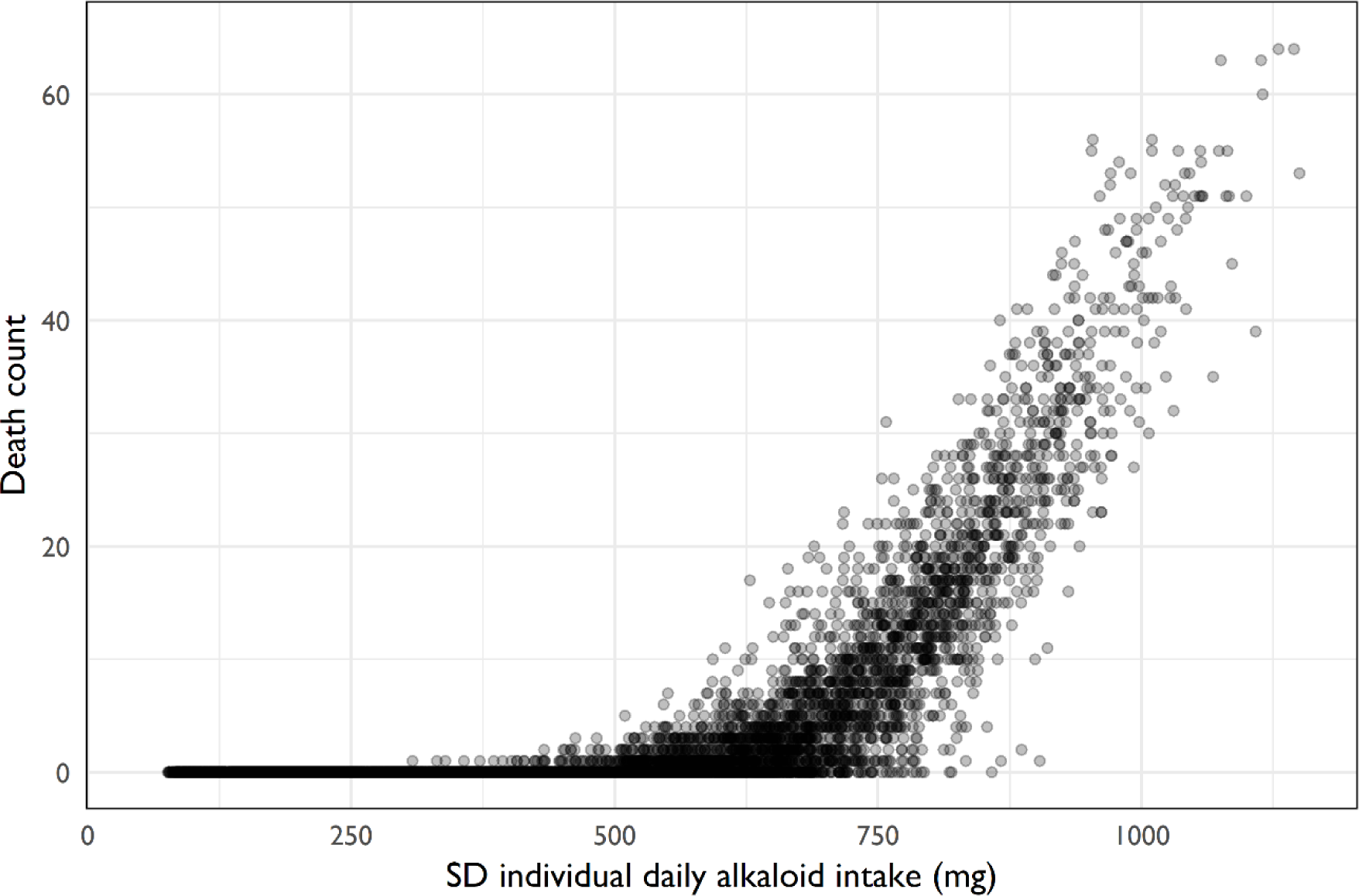
Distribution of model-run death count by model-run standard deviation of individual daily alkaloid intake (mg) (n=7200).

### Larkspur patchiness and forage overlap

Larkspur patchiness exerted a strong influence on intra-herd variation in alkaloid consumption and thus deaths (Fig 4). Total deaths for different levels of LPF ranged from 0 (LPF=0, n=900 model runs) to 13,057 (LPF=5, n=900), with a threshold evident at LPF=3. An examination of the relationships between landscape metrics and deaths using a global linear model and comparison of AICc scores indicated that the model containing only largest patch index was best (AICc=141.7). All other model combinations had ΔAICc values of at least 7.93, indicating all were much less plausible models, given the data (Anderson 2008). The next best univariate model contained only the intercept.

**Fig 4.**
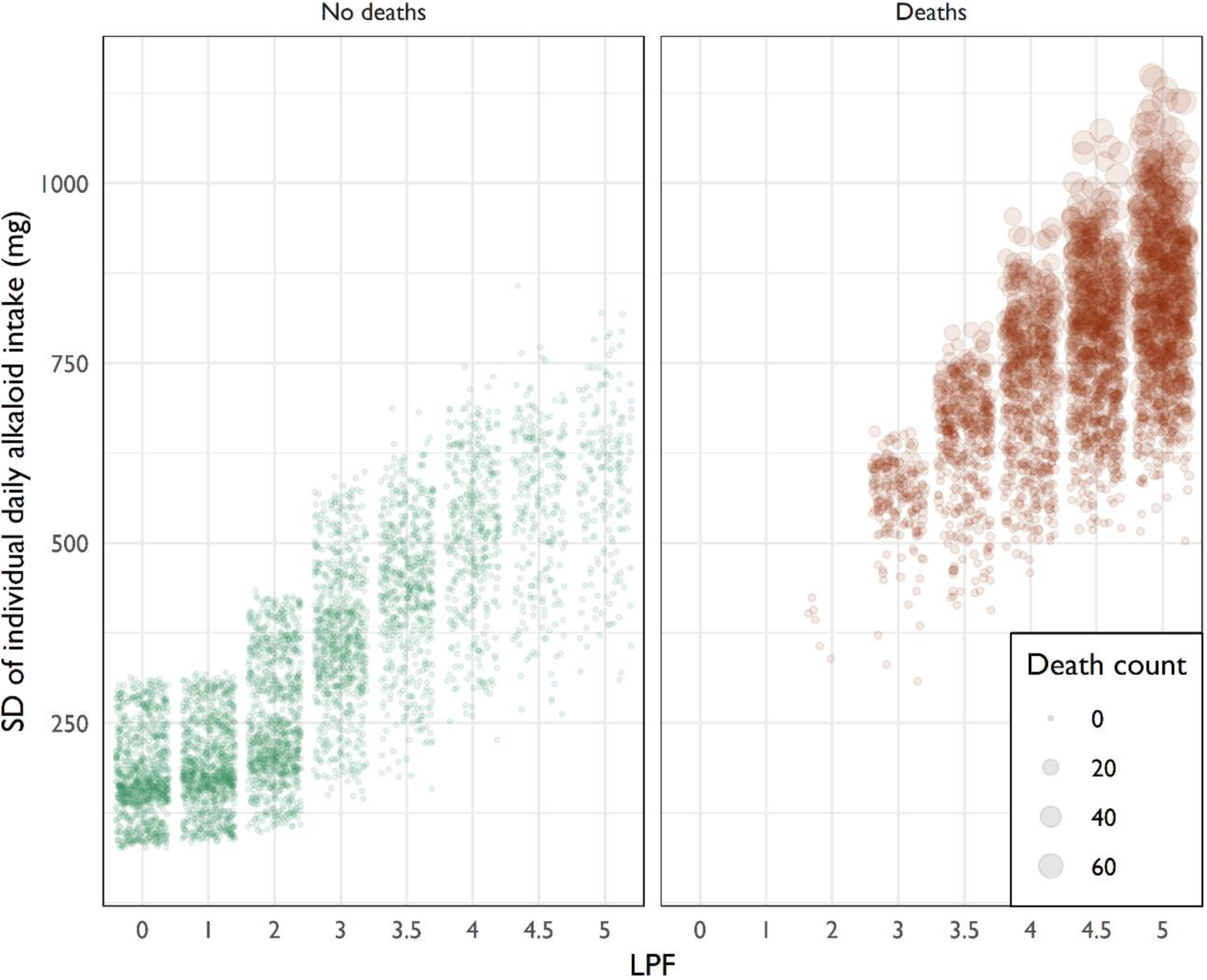
Distribution of model-run standard deviation of individual daily alkaloid intake (mg) by larkspur-patch-factor (LPF) across all levels of other variables. For visibility, data are split by whether or not any deaths occurred during the model run, with points sized to indicate the number of deaths. Points are semi-transparent so that darker areas indicate more points (n=7,200).

Deaths were distributed more evenly among the different levels of larkspur-forage overlap than among the levels of larkspur patchiness, though there was a peak when there was desirable forage both inside and outside of larkspur patches (LFOF=1-2). Total deaths (Table 3) ranged from a minimum of 1,853 (LFOF=0, n=900) to a maximum of 7,230 (LFOF=1, n=900). Model-run standard deviation of daily alkaloid intake largely mirrored deaths, while mean daily alkaloid intake increased with increasing larkspur-forage overlap.

**Table 3.** Data for total deaths, mean deaths, standard deviation of individual daily alkaloid intake (mg), and mean individual daily alkaloid intake (mg) for different levels of larkspur-forage-overlap (LFOF) across all levels of other variables (n=7,200).

| LFOF | Total deaths | Mean deaths | SD intake | Mean intake |
| --- | --- | --- | --- | --- |
| 0 | 1853 | 1.29 | 392.31 | 337.76 |
| 1 | 7230 | 5.02 | 554.64 | 524.98 |
| 2 | 6890 | 4.79 | 563.80 | 575.85 |
| 3 | 6324 | 4.40 | 552.92 | 617.70 |
| 4 | 5440 | 3.78 | 535.08 | 624.47 |

Additionally, there were distinctly different relationships among mean alkaloid intake and the standard deviation of alkaloid intake at low, medium, and high LFOF. With zero larkspur-forage overlap, an increase in alkaloid intake within a model run usually led to increased variation in intake among the herd, leading to increased deaths (Fig 5). When there was high overlap between forage and larkspur, increases in alkaloid consumption within the herd usually led to decreased standard deviation, reducing deaths. At moderate levels (LFOF=1-2), this relationship was more muddled. Each of these effects was modified by larkspur patchiness in a complex interplay illustrated by Fig 5.

**Fig 5.**
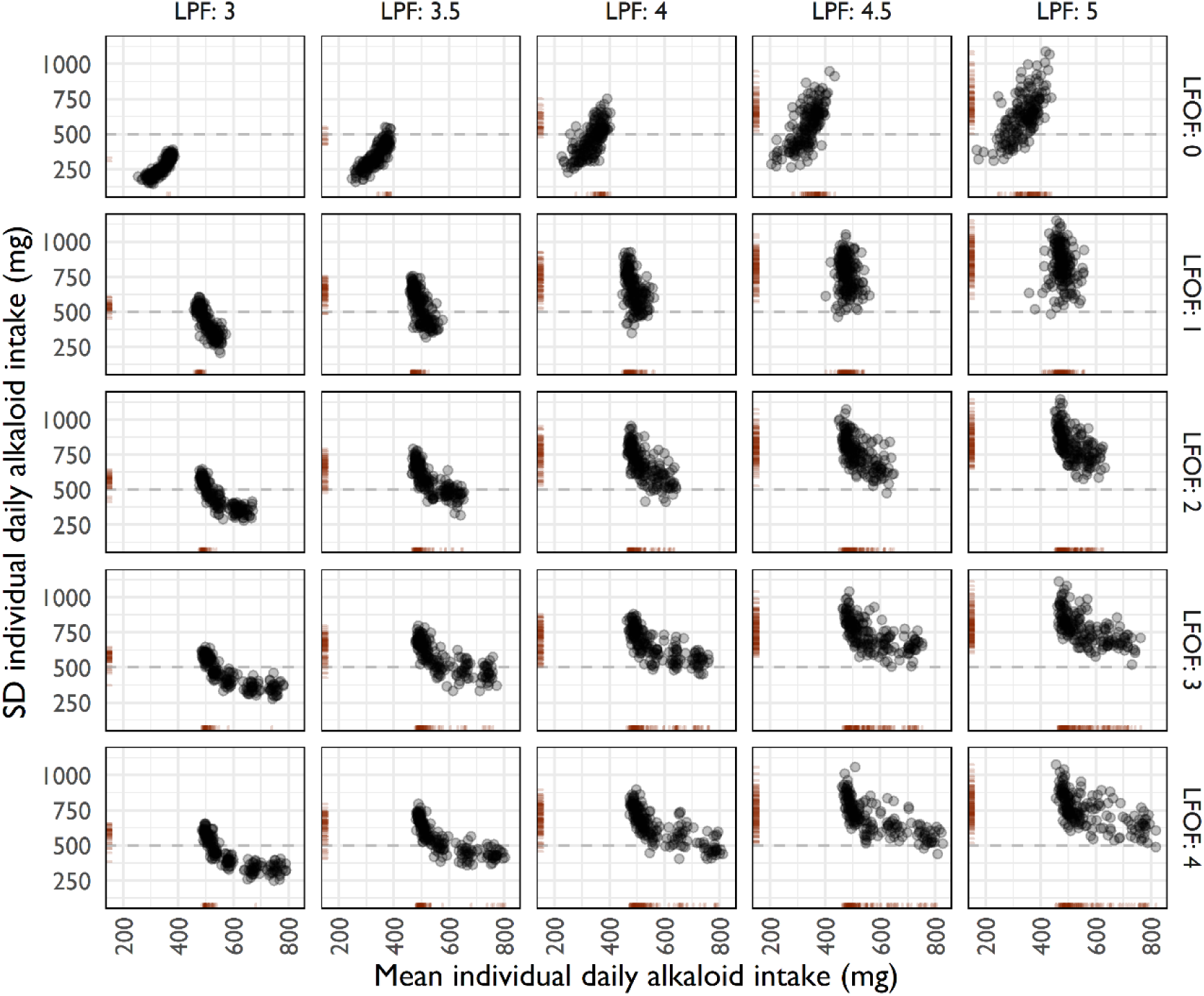
The relationship between mean individual daily alkaloid intake (mg) and standard deviation of individual daily alkaloid intake (mg) at different levels of larkspur-patch-factor (LPF) and larkspur-forage-overlap-factor (LFOF), across all levels of herd-distance-factor. Displayed results are limited to LPF≥3, where the vast majority of deaths occurred. Rug plots on the x and y axes show the distribution of deaths. A dashed line marks a standard deviation of 500 mg, an apparent threshold where deaths increase greatly. Points are semi-transparent so that darker areas indicate more points (n=7,200).

### Herd cohesion

Inter-animal distance was an important factor in alkaloid toxicosis deaths. Regardless of larkspur patchiness and larkspur-forage overlap, just 14.4% of model runs at the minimum herd distance level (HDF=0.5) had at least one death, while 56.3% of model runs resulted in at least one death at the maximum herd distance level (HDF=16). Overall, mean deaths per model run ranged from 0.72 at HDF=0.5 to 8.67 at HDF=16.

The relationship between patchiness, overlap, and herd behavior becomes clearer when larkspur patchiness, larkspur-forage-overlap, and herd distance are used to plot standard deviation of alkaloid consumption and total deaths (Fig 6). Increases in herd distance consistently generated increases in variation in alkaloid consumption, with the magnitude modified by larkspur patchiness and larkspur-forage overlap. However, deaths did not begin to occur until standard deviation approached the threshold of 500 mg, with this being reached at different levels depending on herd cohesiveness and plant patchiness. This means that the degree of herd cohesiveness necessary to prevent deaths was determined by the patchiness of the threat.

**Fig 6.**
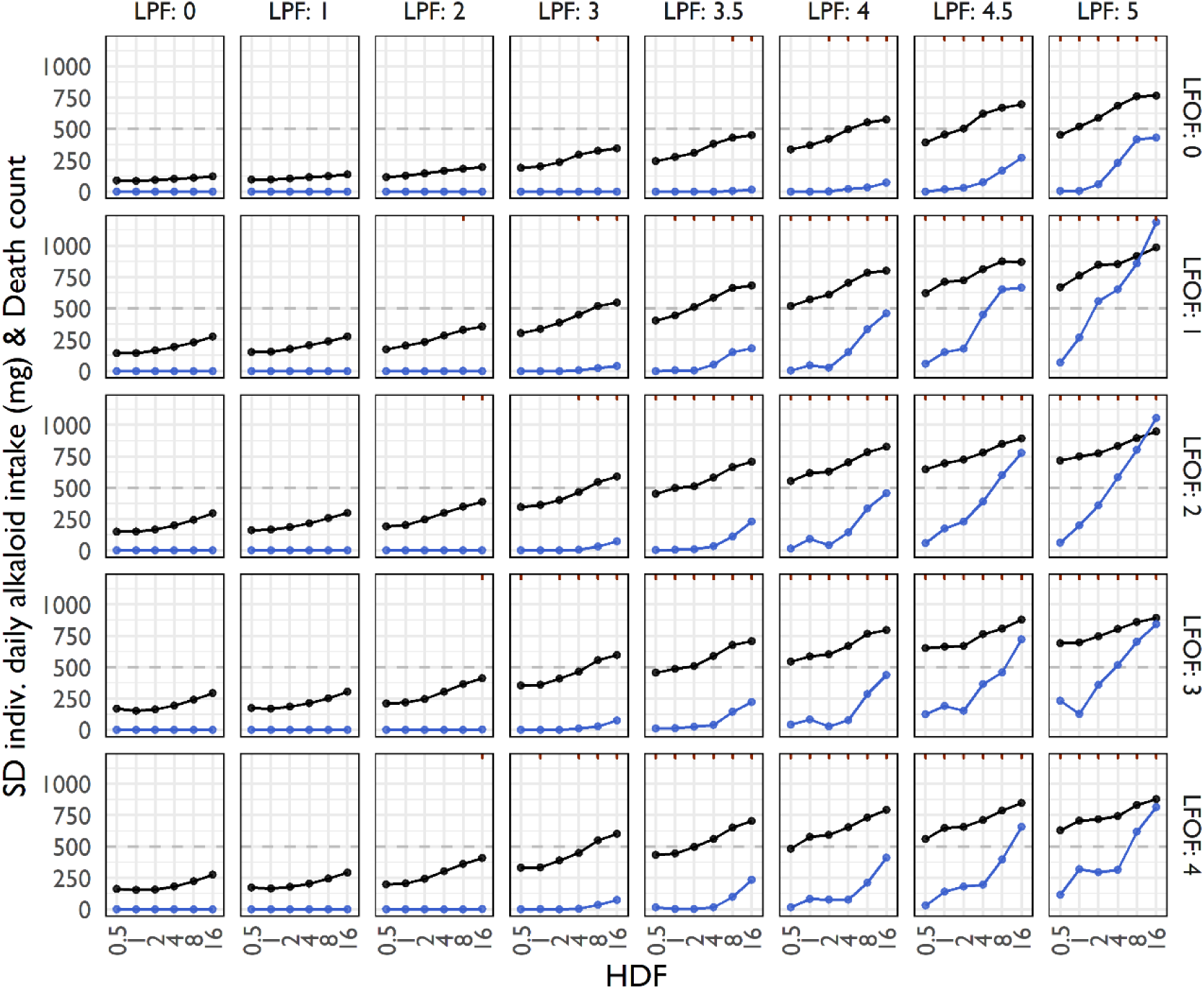
The relationship of herd-distance-factor (HDF), larkspur-patch-factor (LPF), and larkspur-forage-overlap (LFOF) to standard deviation of individual daily alkaloid intake (black) and total deaths (blue). Hash marks on the upper x axis indicate levels where at least one death occurred, and a dashed horizontal line marks a standard deviation of 500 mg, an apparent threshold where deaths begin to occur. Points represent mean model-run values (n=30 per point, 7,200 overall).

### 1/N and encounter dilution

The relationship of “plant predators” to the 1/N concept of predation risk reduction in herds is best understood at LPF=5 and LFOF=4, where there was one large and dangerous patch that overlapped with highly desirable forage, meaning that encounter was inevitable. If we restrict the analysis to only those days when at least one cow consumed larkspur, we can see the distribution of risk when encounter occurred (Fig 7).

**Fig 7.**
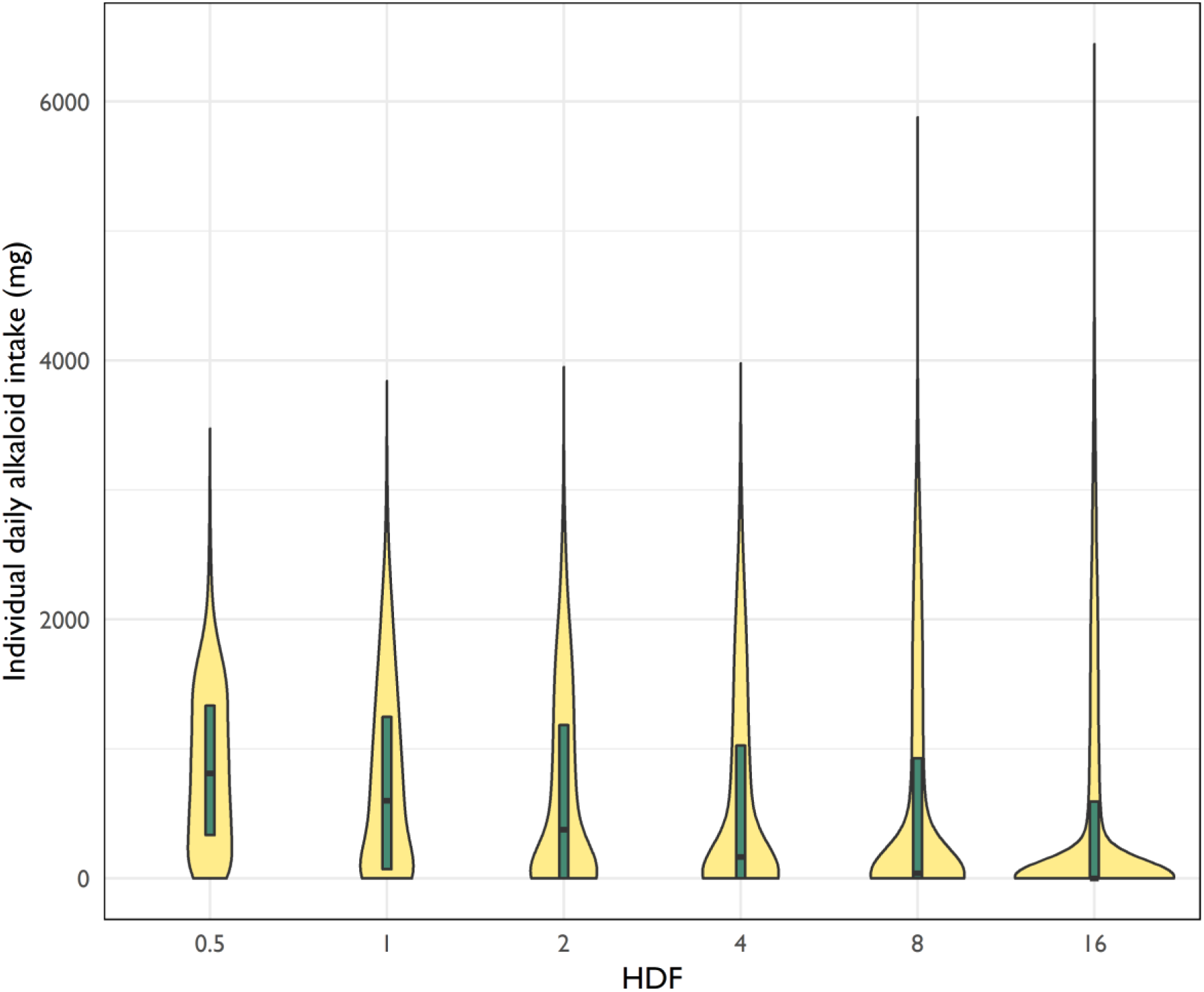
Violin plots of the distribution of individual daily alkaloid intake at different levels of herd-distance-factor (HDF), restricted to highly patchy larkspur overlapping completely with highly desirable forage (LPF=5, LFOF=4) and days where at least one cow consumed larkspur (n=167,383). Within the violins, box plots show the location of the median and first and third quantiles.

In herds with high inter-animal distance (e.g., HDF=16) many cows avoided larkspur encounter entirely, while others consumed a great deal of larkspur, thereby dying. On the other hand, in herds with low inter-animal distance (e.g., HDF=0.5) few cows avoided larkspur entirely, with consumption concentrated at sub-lethal levels. In other words, in highly cohesive herds encountering a serious threat, when one cow encountered larkspur it was likely that all cows in the herd would, reducing the distribution of individual risk and resulting in fewer deaths.

Encounter dilution, where cohesive herds avoid detection by predators with limited capacity to find them, is best understood at LPF=5 and LFOF=0. In this circumstance, a single larkspur patch is undesirable for foraging but a serious threat to cows that nevertheless encounter it. Table 4 shows rates of larkspur encounter and death among different levels of HDF under these conditions. For herds grazing at HDF=0.5, 38.2% of grazing-days passed without a single animal encountering larkspur. On the other hand, herds grazing at HDF=16 managed to entirely avoid larkspur on just 9.4% of grazing-days. This contributed to substantially different rates of death occurrence.

**Table 4.** Percent of grazing-days where larkspur was encountered or a death occurred, at different levels of herd-distance-factor (HDF). Data are restricted to cases where highly patchy larkspur did not overlap at all with highly desirable forage (LPF=5, LFOF=0) (n=3,240).

| HDF | Larkspur encountered? |  | Death occurred? |  |
| --- | --- | --- | --- | --- |
|  | No | Yes | No | Yes |
| 0.5 | 38.2% | 61.9% | 99.8% | 0.2% |
| 1 | 33.7% | 66.3% | 98.9% | 1.1% |
| 2 | 31.5% | 68.5% | 97.4% | 2.6% |
| 4 | 24.4% | 75.6% | 93.3% | 6.7% |
| 8 | 22.8% | 77.2% | 85.9% | 14.1% |
| 16 | 9.4% | 90.6% | 79.1% | 20.9% |

## Discussion

Interactions between domestic herbivores and forage plants are complex, with many important spatiotemporal scales of interaction (Wiens 1976, Launchbaugh and Howery 2005, Larson-Praplan et al. 2015). Perhaps due to the relative simplicity, most research attention has been paid to the interaction of individual livestock with individual plants (including sequences of individuals), and the consequent effects on the grazer and the grazed (e.g., Provenza et al. 2003, Diaz et al. 2007, Villalba et al. 2015). This has been especially true of research on the effect of plant toxins on livestock (Knight and Walter 2001, Welch et al. 2015a). Less common has been research examining aggregations of plants, groups of herbivores, or both. What research there has been in this category has focused largely on how livestock affect plants (e.g., Milchunas et al. 1988, Maschinski and Whitham 1989, Cromsigt and Olff 2008).

Rarest of all has been research seeking to understand how plant patchiness influences group behaviors and outcomes in livestock (though note the significant body of research on “grazing lawns” that at times includes reciprocal relationships between plants and wild herbivores, e.g., McNaughton 1984). Because this type of research requires integration of environmental and animal data at a wide array of scales, it is difficult to design, conduct, and analyze. Nevertheless, if we are to improve our understanding and management of heterogeneity we must expand our capacity to connect pattern and process to illuminate these multiscale relationships (Fuhlendorf et al. 2012).

Here, we have addressed this challenge via the use of a bottom-up agent-based model, incorporating empirical data and neutral landscape models to provide novel insight into why large herbivores may have evolved to respond to plant patchiness with patchiness of their own. Our results show that herd behavior and plant patchiness interact in a complex but conclusive manner to generate or mitigate risk from dangerous plant toxins, with important implications for grazing management and for theory on group behavior in herbivores.

### Evaluating hypotheses

Every simulated pasture contained 1.13 million mg of MSAL-type alkaloids, enough to provide 282 lethal doses to 500 kg cows, and each pixel was equally likely to be dangerous, regardless of spatial arrangement. We were thus surprised that disaggregated larkspur, distributed randomly or in small patches, caused zero deaths, even when overlapping completely with desirable forage. Regardless of herd cohesion, deaths did not occur regularly until the largest patch exceeded 3,800 m^2^, with 4.3 ha of larkspur divided among 31 patches or fewer. Clearly, patchy larkspur kills, and non-patchy larkspur does not. Despite occasional observations in the literature of the patchy growth of most dangerous larkspur species (Kotliar 1996, Pfister et al. 2010), this is a novel conclusion.

Results for larkspur-forage overlap ran counter to our hypothesis. We had expected that increased forage draw within larkspur patches would always lead to increased deaths. This was not the case. Instead, deaths were maximized when there was some desirable forage within large larkspur patches but most remained outside of larkspur patches. Fig 5 indicates that even though mean larkspur intake is lower in these situations, intake variation among individuals in the herd is higher. Thus, it appears that moderate levels of larkspur-forage overlap effectively split herds, with some individuals entering larkspur patches and others remaining outside to graze other desirable forage.

### Behavioral ecology of herds

The 1/N effect typically describes a situation where a predator can capture one (or whatever the numerator value is) prey, thus the chance of any individual being selected declines with an increasing denominator (N). However, we propose that a more flexible way to understand dilution is as *risk*/N. Here, a predator presents potential prey with a certain amount of risk and individual risk is diluted as N increases. In this case, not only is the amount of risk presented by the predator important, but also the distribution of that risk. Assuming equal vulnerability, if the distribution of risk is such that a given herd member will not equal or exceed the level of risk it would acquire on its own, then herd membership is beneficial to the individual. As opposed to 1/N, which usually assumes that at least one death will occur on encounter, *risk*/N allows for cases where risk is so broadly and evenly distributed that all herd members evade death by virtue of simply being in a group.

If we conceptualize larkspur intake as consumption of risk, it is clear that “plant predators” provide an interesting application of *risk*/N. In Fig 7, where at least one herd member has met the predator, members of tightly cohesive herds accumulate greater median risk but with more even distribution. The herd is thus beneficial to the individual not because it lowers absolute risk, but because it lowers the likelihood of accumulating excessive risk when encountering a predator. If the absolute risk presented by a predator is high enough it can still cause death regardless of herd behavior (as in highly patchy larkspur), but it is less likely to regularly do so when risk is evenly distributed amid a cohesive herd.

As noted by Turner and Pitcher (1986), risk of death upon encounter must be considered along with the chance of first encountering predators that have limited perception. In our study, larkspur-forage overlap, which increases the likelihood of the herd encountering larkspur, was akin to perception, so this phenomenon is best illustrated by limiting overlap, as in Table 4. In these circumstances, it is clear that more cohesive herds are less likely to encounter the threat. This largely holds true at different levels of larkspur-forage overlap, though at high levels of overlap moderate levels of herd cohesion lead to the fewest encounters. Nevertheless, overall death counts (Fig 6), which incorporate the benefits of both *risk*/N and encounter dilution, indicate that tightly cohesive herds provide the best overall strategy for avoiding predation by plant predators.

### Limitations

These results must be considered within the context of other benefits and detriments of herd behavior. For example, within the model, individuals in the most cohesive herds traveled 56% greater distance than individuals in herds with the least cohesion. This may indicate that less cohesiveness is desirable when the threat from larkspur is low, as increased cohesion is likely to increase energy expenditure. However, even in this case this observation is offset by the fact that the most cohesive herds met their forage needs 9% faster than the least cohesive herds, likely due to reaching desirable forage more quickly when traveling to stay with the herd. These are complex phenomena, so simple answers are unlikely.

Ultimately, it is most important to recognize that our model was designed to address the questions analyzed here and was not intended to fully replicate cattle behavior. Notably lacking are the more complex (and poorly understood) elements of inter-animal interactions, such as those mediated by familial relationships. Nevertheless, we are confident that our conclusions are sound within the context of the questions we asked.

### Conclusions and implications

In his influential review of “population responses to patchy environments”, Wiens (1976 p. 97) observed that the “patch structure of resources in space and/or their transiency in time governs the form of social organization expressed within a population…”. Even 42 years later, this strikes us as a bold and insightful statement, as empirical evidence for the influence of resource patchiness on social organization remains rather weak (outside of social insects). This study provides clear evidence that social organization in large herbivores can be an adaptive response to patchily distributed poisonous plants.

However, Wiens (1976 p. 96) also wrote that “[s]ocial patterns have no unitary adaptive function, but are the creations of multiple selective pressures, and are thus likely to confer multiple adaptive advantages to individuals”. Even if herd cohesion mitigates plant toxin risk and this functions similarly to demonstrated mechanisms for predation risk mitigation, we think it is unlikely that herd behavior would emerge from the sole pressure of plant toxins. Instead, as Wiens suggested, a strategy as durable as herd behavior in large herbivores is likely to be an adaptation to many pressures, including predation, mate-finding, and heterogeneous forage resources. Here, we have added poisonous plants to that list.

While the benefits of social grouping are well documented in wild herbivores, they have been largely ignored in domestic herbivores, especially within production agriculture in the US and Europe. The result is livestock that are ill-prepared to deal with the pressures that herd cohesion mitigates (e.g., Laporte et al. 2010). Having demonstrated that increased herd cohesion alone can reduce larkspur-induced deaths by greater than 90% in a variety of scenarios, we suggest that the time has arrived for managers to reconsider the importance of herd behavior in their cattle. Because the adaptive functions of herds are manifold, it is likely that the benefits of a renewed focus on herd behavior in our domestic livestock will be manifold as well.

## Acknowledgements

We thank Joel Vaad, manager of the Maxwell Ranch, for support and advice. Early conversations with Michael A. Smith and the Sims family of McFadden, WY provided crucial insight into the relationship between grazing management and larkspur. Jasmine Bruno and David Augustine made small suggestions that led to important changes to the model.

